# Modeling gene x environment interactions in PTSD using glucocorticoid-induced transcriptomics in human neurons

**DOI:** 10.1101/2021.03.01.433391

**Authors:** Michael S. Breen, Tom Rusielewicz, Heather N. Bader, Carina Seah, Changxin Xu, Christopher J. Hunter, Barry McCarthy, Mitali Chattopadhyay, Frank Desarnaud, Iouri Makotkine, Janine D. Flory, Linda M. Bierer, Migle Staniskyte, NYSCF Global Array Team, Scott A. Noggle, Daniel Paull, Kristen J. Brennand, Rachel Yehuda

## Abstract

Post-traumatic stress disorder (PTSD) results from severe trauma exposure, but the extent to which genetic and epigenetic risk factors impact individual clinical outcomes is unknown. We assessed the impact of genomic differences following glucocorticoid administration by examining the transcriptional profile of human induced pluripotent stem cell (hiPSC)-derived glutamatergic neurons and live cultured peripheral blood mononuclear cells from combat veterans with PTSD (*n*=5) and without PTSD (*n*=5). This parallel examination in baseline and glucocorticoid-treated conditions resolves cell-type specific and diagnosis-dependent elements of stress response, and permits discrimination of gene expression signals associated with PTSD risk from those induced by stress. Computational analyses revealed neuron-specific glucocorticoid-response expression patterns that were enriched for transcriptomic patterns observed in clinical PTSD samples. PTSD-specific signatures, albeit underpowered, accurately stratify veterans with PTSD relative to combat-exposed controls. Overall, *in vitro* PTSD and glucocorticoid response signatures in blood and brain cells represent exciting new platforms with which to test the genetic and epigenetic mechanisms underlying PTSD, identify biomarkers of PTSD risk and onset, and conduct drug-screening to identify novel therapeutics to prevent or ameliorate clinical phenotypes.

## INTRODUCTION

Although unequivocally precipitated by environmental events, only a small percentage of trauma survivors develop post-traumatic stress disorder (PTSD)^1^, prompting a search for risk factors that increase the probability of developing this condition following trauma exposure. Convergent lines of evidence are consistent with a heritable component to PTSD risk. Twin studies have demonstrated a concordance between PTSD or lack thereof, in monozygotic and dizygotic twins,^2,3^ and genome-wide association studies (GWAS) have estimated SNP-based heritability from 5–30%^4–6^. The two most recent GWAS each identified genome-wide significant loci associated with PTSD^4,6^, although loci differed across ancestry, sex, and type of trauma. This highlights the necessity of developing paradigms that might help examine the contribution of genetic factors in elevating risk for PTSD following exposure to an environmental stressor.

Insights into the gene × environment (G×E) interactions underlying the psychiatric symptoms of PTSD remain poorly resolved, largely due to the lack of a cohesive neurobiological framework to investigate these mechanisms. While investigations of patient postmortem brain tissue are scarce, recent work suggests that PTSD symptoms may ultimately reflect dysfunction of excitatory neurons^7^, arising as a result of an aberrant or exaggerated stress responses. However, to date, most studies of PTSD pathophysiology have focused on blood cells, the most notable of which have involved stimulation with glucocorticoids, with PTSD patients showing an exaggerated lymphocyte sensitivity to glucocorticoids relative to trauma-exposed controls^8^. Indeed, we recently reported evidence of reduced glucocorticoid response in cultured peripheral blood mononuclear cells (PBMCs) from combat-exposed veterans with PTSD relative to combat-exposed controls^9^. Thus, in order to resolve the relative contributions of genetic risk and environmental stress to PTSD pathophysiology, it is critical to deconvolve the impact of stress in a cell-specific manner, including determinants of stress responses across brain and blood cells. Notably, glucocorticoid receptor signaling in blood and brain are convergent in murine models and highly associated with individual differences in response to trauma exposure^10^. Nevertheless, the degree to which PTSD symptoms arise as a result of elevated/sustained peripheral response to stress^8,10^ and/or abnormal neuronal or circuit response to peripheral stress cues^11–13^ remains unresolved. This work seeks to uncover GxE interactions between heritable genetic and/or epigenetic factors with stress exposure utilizing PTSD patient-derived human neurons.

An improved understanding of the pathophysiology of PTSD requires the development of appropriate human-specific models. Understanding the extent to which the dysregulated stress response reflects cell-type-specific preexisting genetic vulnerabilities will improve genetic-based diagnosis and potentially identify novel therapeutic targets to prevent or reverse PTSD. Human induced pluripotent stem cell (hiPSC)-based models have the potential to address some of the major limitations of clinical and animal-based studies by generating human neural cells that are genetically identical to those found in patients. Although there have been a handful of studies of glucocorticoid response in live human neurons^14,15^, none have contrasted neurons derived from PTSD cases and trauma-exposed controls.

Here we describe the first hiPSC-based study of PTSD, comprised of five veterans with PTSD and five combat-exposed controls. Although we did not detect baseline differences between PTSD and control-derived blood or neurons, we identified diagnosis-dependent differences in response to glucocorticoid treatment. Using matched PBMCs from this cohort, and validated in a larger independent blood cohort^9^, we describe shared and cell-type-specific aspects of stress response. Overall, given the fetal-like nature of hiPSC-derived neurons^16^, we conclude that critical aspects of glucocorticoid response are encoded in patient genetics, consistent with a clear genetic predisposition to PTSD. By coupling analysis of glucocorticoid response in patient-matched blood and hiPSC-neurons, we report that straightforward analysis of blood biomarkers can be used to predict PTSD with high accuracy. The current work indicates that if the biological mediators of environmental risk can be predicted, then hiPSC-based models can be used to test genotype-dependent and cell-type-specific environmental responses. This makes possible causal conclusions as to whether long-standing blood biomarkers represent a cause or consequence of disease, and hint at possible therapeutic strategies to minimize the likelihood of PTSD following trauma exposure, a particularly valuable outcome for military and first-responder personnel.

## MATERIALS AND METHODS

### Participants

Participants were combat-exposed veterans with and without PTSD (*n* = 5, respectively) who provided written, informed consent and for whom a viable fibroblast sample was provided and sufficient RNA for genome-wide expression analyses was extracted. Eligibility for participation was determined as previously described^9^. In brief, all participants underwent psychological evaluation using the Structured Clinical Interview for DSM-5 (SCID) and the Clinician Administered PTSD Scale (CAPS) for determination of PTSD diagnosis and severity. Diagnostic and clinical exclusions included: i) presence of moderate or severe substance use disorder within the past 6 months, ii) lifetime history of primary psychotic or Bipolar I disorders, iv) neurological disorder or major systemic illness, and v) treatment with systemic steroids; for PTSD–, vi) current or recurrent major depressive disorder were exclusionary.

### Biopsy Collection and generation of hiPSCs

Fibroblasts from PTSD and combat-exposed control subjects were isolated at the James Peter VAMC, per Institutional Review Board–approved human subject protocol. Skin fibroblasts of PTSD and non-PTSD patients were collected via skin punch biopsy from an area of the body at the doctor’s discretion. Upon collection, biopsies were immediately placed in Biopsy Collection Media (Cascade Biologics Medium 106 (Life Technologies, M-106-500), 1X Antibiotic-Antimycotic (Life Technologies, 15240-062), LSGS (Life Technologies, S-003-10)) and stored at 4 °C for a maximum of 24 hours. A courier was then used to transport the biopsies in Biopsy Collection Media to the New York Stem Cell Foundation, where they were reprogrammed into hiPSCs as previously described using the NYSCF Global Stem Cell Array®^17^.

In brief, biopsies were dissected into ~1 mm^3^ pieces and plated in Biopsy Plating Media (Knockout-DMEM (Life Technologies, 10829-018), 10% FBS (Life Technologies, 100821-147), 2 mM GlutaMAX (Life Technologies, 35050-061), 0.1 mM MEM Non-Essential Amino Acids (Life Technologies, 11140-050), 1X Antibiotic-Antimycotic (Life Technologies, 15240-062), 0.1 mM 2-Mercaptoethanol (Life Technologies, 21985-023)). Upon the outgrowth of fibroblasts, samples were entered into an automated pipeline producing vials of cells for both iPSC reprogramming and backup stock. A cell pellet was collected, with DNA isolated using an EPMotion and the ReliaPrepTM 96 gDNA Miniprep HT System (A2671, Promega). This DNA was used to confirm a sample’s chain of custody throughout the reprogramming process. Fibroblasts were reprogrammed using modified mRNA (Reprocell, 000076) and enriched using anti-fibroblast meads (Miltentyi Biotec, 130-050-601) in automated procedures.

Reprogrammed iPSCs were then expanded using PSC Feeder Free Media (Thermo Fisher Scientific: A14577SA) and grown on Cultrex coated plates (R&D Systems, 3434-010-02). Cells were routinely passaged using our automated platform in the presence of Accutase (ThermoFisher, A11105-01). Following passage cells were maintained in PSC media supplements with 1 μM thiazovivin (Sigma-Aldrich, SML1045-25MG). All cells were frozen in Synthafreeze (ThermoFisher, A12542-01) into master stocks before being QC’d. As part of the QC process, all samples were tested for Mycoplasma (Lonza, LT07-710) and Sterility (Hardy Diagnostics, K82), alongside a confirmation of karyotypic stability (Illumina, 20030770), post-thaw viability recovery (greater than 50% confluency reached within 10 days), correct donor identity (Fluidigm, SNPTrace), iPSC gene expression (Nanostring, Custom Panel) and a test for differentiation potential (Custom Panel, Nanostring) as previously described^18^.

### Generation of hiPSC-derived mixed forebrain neurons and incubation with hydrocortisone

The hiPSCs were cultured on 6-well tissue culture plates coated with Matrigel in 5 mL of DMEM/F12 medium (1:100 dilution), and the cells were routinely passaged after reaching 80% confluence or colonies are ready for neural differentiation via an embryoid body (EB). EBs were grown for 7-10 days. Followed by plating EBs to Poly-L-Ornithine/Laminin coated plates. Within a week visible rosette began to appear. Then the neural rosettes were enzymatically collected after 14 days of culture by Rosette Selection Reagent (StemCell Technologies) and plated into matrigel-coated plates. Resulting neural progenitor cells (NPCs) were maintained on Matrigel-coated plates in NPC media (DMEM/F12 supplemented with 1x N2, 1x B27 (both ThermoFisher Scientific), 1% Antibiotic-Antimycotic and 20ng/ml recombinant fibroblast growth factor 2 (R&D Systems)) and passaged using Accutase (Innovative Cell Technologies)^19^. Magnetic cell sorting was carried out using the cell surface marker CD133 (Promonin 1), similar to Goate *et al*., 2019, to enrich for the expandable NPCs. Briefly, magnetic activated cell sorting (MACS) was carried out following a two-step protocol, beginning with CD271 depletion, followed by CD133 selection^20^. NPC lines were immunohistochemically validated using antibodies for NESTIN. After the validation, NPCs were expanded in NPC media and differentiated for six weeks into mixed FB neurons. After six weeks, mixed FB neurons were treated with hydrocortisone (0.1 μM, 1 μM and 2.5 μM) for 24 hours.

### Generation of hiPSC-derived NGN2-neurons and incubation with hydrocortisone

hiPSCs were differentiated to induced glutamatergic neurons as previously described with some modifications^21^. hiPSCs were single cell passaged after a 20-minute dissociation with Accutase (STEMCELL Technologies) at 37 °C and 5% CO2. 1 million cells per well were plated in 12-well Cultrex coated (R&D Systems, 3434-010-02) tissue culture plates (Corning Costar) in PSC Feeder Free Media (Thermo Fisher Scientific: A14577SA) with 1 μM thiazovivin (Sigma-Aldrich: SML1045). Lentivirus (generated by ALSTEM) carrying pLV-TetO-hNGN2-eGFP-Puro (Addgene#79823) and FUdeltaGW-rtTA (Addgene#19780) was diluted to an MOI=1 each (1 million genome counts of each vector per transduction) in 100 μL DPBS, no calcium, no magnesium (Thermo Fisher Scientific) and added directly after cell seeding. After 24 hours, the medium was exchanged with Neural Induction Medium (NIM) comprising a 50:50 mix of DMEM/F12 and Neurobasal, with 1X B27+Vit.A, 1X N2, 1X Glutamax (Thermo Fisher Scientific), and 1 μM doxycycline hyclate (Sigma-Aldrich). 24 hours later the medium was removed and NIM was added with doxycycline plus 5 μg/mL puromycin. A full medium exchange was performed the next day with NIM plus doxycycline and puromycin. 24 hours later cells were passaged by incubating with Accutase for 30 minutes at 37 °C and 5% CO2. 96-well plates (PerkinElmer CellCarrier Ultra) were coated with 0.1% Polyethylenimine (PEI) (Sigma-Aldrich: 408727) in 0.1 M Borate buffer pH 8.4 for 30 minutes at room temperature, washed 5x with water and prefilled with 100 uL per well of Neural Coating Medium (NCM) comprising of Brainphys medium (STEMCELL Technologies) with 1X B27+Vit.A, 1 μM thiazovivin, 5 μg/mL puromycin, 250 μM dbcAMP (Sigma-Aldrich), 40 ng/mL BDNF (R&D Systems), 40 ng/mL GDNF (R&D Systems), 200 μM ascorbic acid (Sigma-Aldrich) and 10 μg/mL natural mouse laminin (Sigma-Aldrich). A sample of cells were stained with 10 μg/mL Hoechst μLus 1:500 acridine orange/propidium iodide solution and counted on a Celigo imager (Nexcelom Bioscience). 50,000 cells per well were seeded into NCM filled 96-well plates in 100uL per well of Neural Medium (NM) comprising of Brainphys medium with 1X B27+Vit.A, 1 μM thiazovivin, 5 μg/mL puromycin, 250 μM dbcAMP, 40 ng/mL BDNF, 40 ng/mL GDNF, 200 μM ascorbic acid and 1 μg/mL natural mouse laminin. 24 hours after seeding medium was exchanged to Neural Selection Medium (NSM) comprising of Brainphys medium with 1X B27+Vit.A, 250 μM dbcAMP, 40 ng/mL BDNF, 40 ng/mL GDNF, 200 μM ascorbic acid, 1 μg/mL natural mouse laminin and 2 μM Arabinosylcytosine (Sigma-Aldrich). 48 hours later NSM medium was fully exchanged. 48 hours after the medium was fully exchanged with Neural Maintenance Medium (NMM) comprising Brainphys medium with 1X B27+Vit.A, 250 μM dbcAMP, 40 ng/mL BDNF, 40 ng/mL GDNF, 200 μM ascorbic acid and 1 μg/mL natural mouse laminin. Thereafter every 48 hours, half the medium was exchanged with NMM until day 21 post transduction passage. All medium exchanges were performed using a Hamilton Star liquid handler set to 5 μL/second for aspirate and dispense as part of the NYSCF Global Stem Cell Array®^17^.

At day 21, HCort treatment medium was prepared by first dissolving HCort (Sigma-Aldrich: H0888) in ethanol to make a 2.8 mM stock. HCort ethanol stock was then diluted to 0.2 mM in HBSS. Final treatment medium was prepared by diluting HCort HBSS stock into NMM to a final concentration of 0.1, 1 and 2.5 μM which was then applied to cells by fully exchanging medium. Ethanol was equalized to 15 μM in control and all treatment media. 24 hours after HCort treatment all medium was removed using the Bluewasher (BlueCatBio) and cells were lysed for 5 minutes using RLT plus buffer (Qiagen), snap frozen on dry-ice and stored at −80 °C. A replicate plate was fixed for immunofluorescence analysis by adding 32% Paraformaldehyde (Electron Microscopy Sciences) directly to medium to a final concentration of 4% and incubated at room temp for 15 minutes. Cells were washed three times with HBSS (Thermo Fisher Scientific), stained overnight with mouse anti-Nestin 1:3000 (Millipore: 09-0024), chicken anti-MAP2 1:3000 (Abcam: 09-0006) in 5% Normal Goat Serum (Jackson ImmunoResearch) in 0.1% Triton x-100 (Thermo Fisher Scientific) in HBSS. Primary antibodies were counterstained with Goat anti-Mouse Alexa Fluor 555 and Goat anti-chicken Alexa Fluor 647 and 10 μg/mL Hoechst for 1 hour at room temp. Cells were washed three times with HBSS and nine 20x fields were imaged per well (2 wells per hiPSC line) using the ArrayScan automated microscope (Thermo Fisher Scientific).

### Optimization of glucocorticoid treatments and concentrations

Preliminary studies were conducted to identify optimal culture and glucocorticoid stimulation conditions. These pilot studies sought to evaluate the transcriptional effects of hydrocortisone (HCort) and dexamethasone (DEX) on mixed FB and NGN2 neurons, optimize the length of glucocorticoid treatment and concentrations, and determine whether NGN2 or mixed FB cell cultures show greater response to glucocorticoid stimulation. These approaches build upon our previously published functional assay, which measures the responsiveness to glucocorticoids^9^. First, we examined the transcriptional perturbations of DEX in neurons by performing a series of qPCR assays. These experiments tested six well-described glucocorticoid regulatory genes across NGN2 neurons and three different neuronal cell lines, covering 11 distinct concentrations of DEX (0 nM, 0.5 nM, 1 nM, 2.5 nM, 5 nM, 10 nM, 27.5 nM, 50 nM, 100 nM, 550 nM, 1100 nM). No distinct gene expression differences were observed across DEX concentrations (**Figure S1**). Second, we sought to optimize conditions for HCort administration to neurons, which, like DEX, is a glucocorticoid agonist with a high affinity to the glucocorticoid receptor. Importantly, HCort has previously proven effective in creating strong glucocorticoid-induced expression changes in neurons. We also rationalized that examination of a mixed population of neurons (*i.e*. mixed FB neurons) might respond differently. Thus, we carried out another set of pilot studies, this time generating genome-wide RNA-seq (*n*=3 donors), to examine the effects of 1000nM HCort (relative to vehicle condition) on NGN2 and mixedFB neurons following a 6 hour and 24 hour incubation period. Based on our pilot work with DEX, we hypothesized that 1000nM of HCort would produce a large genome-wide effect. These results illustrate that 24 hour incubation produced a strong genome-wide effect among a small number of genes, with many other candidate genes trending in their significance. Thus, for the final selection of HCort dosages in the current study, we included a vehicle untreated condition (0 nM), along with a low dose (100 nM), medium dose (1000 nM) and a high dose (2500 nM).

### PBMC isolation and dexamethasone treatment

We followed identical procedures for isolation of PBMCs, cell culture methods and incubation with DEX as recently described^9^. In brief, blood was collected in ethylenediaminetetraacetic acid (EDTA)-containing vacutainer tubes (VWR, West Chester, Pennsylvania) and PBMCs were isolated by density gradient centrifugation using Ficoll-Paque (GE Healthcare) and washed twice in HBSS. Mononuclear cells were then counted manually using a Cellometer Disposable Counting Chambers (Nexcelom Bioscience LLC. Lawrence, MA). The cells were re-suspended in complete RPMI, containing RPMI-1640 (Gibco), 10% fetal bovine serum, 50 U/ml penicillin–streptomycin mixture (Gibco) at a density of 1.75–2.00 × 10^6^ cells/ml of the medium. For in vitro incubation with DEX, 20 μl of DEX at concentrations of 0, 27.5, 55, and 550 nM were pipetted in a flat bottom 96-well plate. To increase RNA yield, a total of 18 wells were prepared for each DEX concentration (~9.0 × 10^6^ cells per dose) in complete RPMI. PBMCs were prepared at 2.5 × 10^6^ cells/ml in complete RPMI and 200 μl was pipetted into each well, bringing the final concentrations of DEX to 0, 2.5, 5, and 50 nM. Following incubation at 37 °C, 5%_(vol/vol)_ CO_2_ for 72 hours, the plates were centrifuged at 900×*g* for 15 min at 4 °C and 160 μl of the supernatant was collected and pooled from each DEX concentration well. The cell pellet on the bottom of each well was re-suspended in 100 μl of TRIzol reagent. Cell lysates for each DEX dose were pooled, aliquoted and stored at −80 °C until RNA isolation.

### RNA extraction and quality control

RNA extraction was performed for mixed FB neurons, *NGN2*-neurons and PBMCs. For mixed FB neurons, following HCort treatment, neurons were collected and stored in TRIzol and RNA extraction was performed with miRNAeasy Mini Kit (Qiagen). For *NGN2*-neurons, RNA was harvested with RNeasy plus micro kit (Qiagen). For PBMCs, RNA was extracted from TRIzol-lysed PBMCs using the miRNAeasy Mini Kit (Qiagen). Following each extraction, RNA quantity was measured on the Nano Drop 2000 Spectrophotometer (Thermo Scientific) and the quality and integrity measured with the Agilent 2100 Bioanalyzer (Agilent, Santa Clara, CA, USA). All RNA integrity numbers (RINs) were high in the current study: mixed FB neurons (9.7 ± 0.39), *NGN2*-neurons (8.8 ± 0.53), PBMCs (7.5 ± 0.95).

### RNA-sequencing data generation

We adopted a low-input RNA-sequencing protocol for the generation of RNA-sequencing data from mixed FB and *NGN2*-neurons. Specifically, polyA enriched RNAs were subjected to RNA-sequencing library preparation using the SMART-Seq v4 Kit (SSv4; Takara) and sequenced using a paired-end 150bp configuration with 30M supporting reads per sample. With regards to RNA-seq data generation from PBMCs, Ribo-zero rRNA deleted RNAs were subjected to RNA-sequencing library preparation using the Illumina TruSeq Stranded Total RNA kit (Illumina) and sequenced using a paired-end 150bp configuration with 20M supporting reads per sample.

### RNA-sequencing data pre-processing

All RNA-sequencing FASTQ files underwent matching analytical procedures, as described previously. In brief, resulting short reads with Illumina adapters were trimmed and low-quality reads were filtered using TrimGalore^22^ (*--illumina* option). All high-quality reads were then processed for alignment using the hg38 reference and the ultrafast universal RNA-seq aligner STAR with default parameters^23^. Mapped bam files were sorted using Samtools and short read data were quantified using featureCounts^24^ with the following parameters: -T 5, -t exon, and -g gene_id. Subsequently, all read counts were exported and all downstream analyses were performed in the R statistical computing environment. Raw count data was subjected to non-specific filtering to remove low-expressed genes that did not meet the requirement of a minimum of two counts per million (cpm) in at least ~ 40% of samples. This filtering threshold was applied to neurons (mixed FB and *NGN2*) and PBMCs separately. All expression values were converted to log_2_ RPKM and subjected to unsupervised principal component analysis (PCA) to identify and remove outlier samples that lay outside 95% confidence intervals from the grand averages. No extreme outliers were detected in the current data sets.

### Developmental specificity analysis

We integrated several RNA-seq datasets from existing postmortem brain tissue and hiPSC models to validate the developmental specificity of our samples using a previously described approach^25,26^. In brief, a total of 16 independent studies were collected covering 2716 independent samples and 12,140 genes. These samples cover a broad range of early hiPSCs, NPSc, mature neuronal cultures, prenatal and postnatal brain tissues. All expression values were converted to log_2_ RPKM and collectively normalized using quantile normalization using the *limma* R package. Subsequently, for each independent sample present in our hiPSC neuronal data set, we performed pair-wise correlation analysis (using Pearson’s correlation coefficients) across all independent samples and subsequently aggregated the correlation coefficients for each external study and/or cell type. Next, and as a complimentary approach, all datasets were jointly analyzed and integrated using principal component analysis (PCA).

### Differential gene expression analyses

All gene expression values were normalized using VOOM normalization^27^ and these data were used to carry out the remainder of downstream analyses. Prior to testing for differentially expressed genes, we applied linear mixed effect models to decompose the transcriptome variability into discrete percentages of variability attributable to multiple known biological and technical sources of variation using the R package variancePartition^28^. Thus, for each gene, the percentage of gene expression variation attributable to differences in induced cell types (*i.e*., hiPSC-mixed FB versus hiPSC-*NGN2*-neurons), individual as a repeated measure (i.e., inter-donor effects), PTSD diagnosis and RIN. As a result, it is possible to identify and partially correct for some confounding variables in our differential gene expression analysis. Next, differential gene expression was conducted using a moderated *t* test from the R package *limma*^27^. PTSD-intendent modules adjusted for the possible confounding influence of PSTD diagnosis and RIN, while PTSD-dependent models factor diagnosis as a main outcome. Due to the repeated measures study design, where individuals are represented by multiple independent technical replicates, the duplicateCorrelation function was applied in the *limma* analysis and gene level significance values were adjusted for multiple testing using the Benjamini and Hochberg method to control the false discovery rate (FDR). Genes passing a FDR <5% were labeled as significantly differentially expressed.

### Weighted gene co-expression network analysis and functional annotation

Signed co-expression networks were built for hiPSC mixed FB and *NGN2*-neurons using weighted gene co-expression network analysis (WGCNA)^29^. To construct a global weighted network for each cell type, a total of 16,933 post QC genes were used. The absolute values of Biweight midcorrelation coefficients (optimal for small sample sizes) were calculated for all possible gene pairs within each cell type and resulting values were transformed using a β-power (β = 12 for mixed FB neurons; β = 7 for *NGN2*-neurons) so that the final correlation matrices followed an approximate scale-free topology. In order to determine which modules, and corresponding biological processes, were most associated with HCort, we ran singular value decomposition of each module’s expression matrix and used the resulting module eigengene (ME), equivalent to the first principal component, to represent the overall expression profiles for each module. Each module was enriched for Gene Ontology (GO) biological processes, molecular factors, cellular components and molecular pathways using ToppGene^30^. All genes passing non-specific filtering in the current data set were used as a genomic background. Only gene sets that passed a multiple test adjustment using the Benjamini Hochberg procedure (Adj. *P* < 0.05) were deemed significant. Notably, no co-expression modules in mixed FB neurons were significantly associated with increasing concentrations of HCort, and thus the current analysis specifically focused on *NGN2*-neurons.

### Gene co-expression module preservation analysis

To identify gene co-expression modules that were either disrupted or created in response to glucocorticoids across *NGN2*-neurons and PBMCs, a permutation-based preservation statistic (Z_summary_) was implemented^31^ as previously described^9^. A preservation statistic (Z_summary_) with 200 random permutations was used to measure the (dis)similarity in correlation patterns for the genes within these gene sets, whereby Z_summary_ > 10 indicates strong evidence of preservation, 2 < Z_summary_ < 10 indicates weak-to-moderate evidence of preservation and Z_summary_ < 2 indicates minimal-to-no evidence of preservation. For this analysis, we specifically focused on dynamically regulated, glucocorticoid responsive functional modules that were identified in either *NGN2*-neurons or PBMCs, respectively.

### Sample size estimates

The R package (ssize.fdr) as proposed by Orr and Liu^32^ was used to calculate the sample size for gene expression. At the FDR of 5%, the proportion of non-differentially expressed genes was computed for each comparison and sample size estimate (and thus varied from 95-99.9%), the desired fold change was fixed at 1.5, the standard deviation was fixed at 0.5, and the desired power at 80%. All power analyses were performed on *VOOM* normalized data.

### Integration of disease-associated genes and gene-sets

Transcriptome-wide summary statics were collected for postmortem brain tissue of individuals with PTSD, major depressive disorder, schizophrenia, bipolar disorder, autism spectrum disorder, alcohol use disorder and inflammatory bowel disease as negative controls. We also collected transcriptome-wide summary statistics from PBMCs of individuals with PTSD, childhood trauma, autism spectrum disorder, major depressive disorder, schizophrenia and bipolar disorder. Genome-wide concordance was examined using Spearman’s correlation coefficient on log_2_ fold-changes. Further, to examine glucocorticoid dysregulation of neurodevelopmental genes we collected genetic risk loci associated with epilepsy, developmental delay, autism spectrum disorder, intellectual disability, schizophrenia, and FMRP target genes. Overrepresentation of genetic risk-related gene sets within differentially expressed gene sets from the current study were analyzed using a one-sided Fisher exact test to assess the statistical significance. All *p*-values, from all gene sets and modules, were adjusted for multiple testing using the Benjamini–Hochberg procedure. An adjusted *p*-value < 0.05 was required to claim that a gene set is enriched within a module. These gene lists (and corresponding references) are available in Supplemental Table 6.

### eQTL enrichment

We examined the overlap between genes whose variance is explained by inter-donor variability and brain cis-eQTLs. To do so, we leveraged the largest collection of eQTLs conducted to date in the human brain together with genes exceeding a variance percentage cutoff for a donor as a repeated measure, as well as other variables of interest in the current study using a previously described technique^26^. In brief, varianceParition analysis assigned each gene a percentage of variance explained by a specific known factor (e.g. donor, RIN). A total of 40 different variance explained cutoff thresholds were examined and the overlap between genes with percentages exceeding this cutoff and the 3000 genes with the smallest *p* values from *cis*-eQTL analysis is evaluated. The overlap is computed for the observed data and 10,000 data sets with the variance percentages randomly permutated. At each cutoff where > 100 genes are represented, the fold enrichment is computed as the observed overlap over the permuted overlap.

## RESULTS

### Hydrocortisone-stimulated transcriptional responses in hiPSC-neurons

To study how glucocorticoids influence gene expression in neurons in PTSD, patient blood samples were reprogrammed into hiPSCs and used to generate neurons from a primary cohort of combat veterans with PTSD+ (*n* = 5) and combat-exposed PTSD-controls (*n* = 5) (**Table 1**). Two distinct populations of hiPSC-derived neurons were examined: induced glutamatergic neurons generated via transient overexpression of the transcription factor *NGN2* (*NGN2*-neurons)^21^ and mixed FB populations comprised of mixtures of glutamatergic neurons, GABAergic neurons and astrocytes and neural progenitors, differentiated by dual SMAD inhibition (mixed FB neurons)^19^ (**Figure S2-S3**). *NGN2*- and mixed FB-neurons were treated with three distinct concentrations of HCort (100 nM, 1000 nM, 2500 nM) and an untreated vehicle condition (0 nM) and subsequently subjected to RNA-sequencing.

**Table 1.** Description of male combat-veterans in the current study.

| **Table 1. Description of male combat-veterans in the current study.** |  |  |  |
| --- | --- | --- | --- |
|  | **PTSD+ (n=5)** | **PTSD- (n=5)** | **P-value** |
| Age (years) | 37.8 (5.36) | 30.6 (4.34) | 4.8E-02 |
| Hispanic, Y/N | 2/3 | 5/0 | 6.9E-02 |
| Race, Caucasian/African American | 3/2 | 3/2 | 1 |
| BDI | 26.8 (7.95) | 5 (4) | 5.9E-04 |
| PCL | 51.4 (7.8) | 9.6 (9.76) | 7.1E-05 |
| Current MDD, Y/N | 0/5 | 0/5 | 1 |
| LT CAPS - intrusion score | 15.2 (3.27) | 1 (1.41) | 9.0E-05 |
| LT CAPS - avoidance score | 6.4 (2.19) | 0.5 (1) | 1.7E-03 |
| LT CAPS - cognition & mood score | 19.6 (4.98) | 3 (3.83) | 9.3E-04 |
| LT CAPS - hyperarousal score | 16.2 (3.9) | 3 (3.56) | 1.2E-03 |
| LT CAPS - total severity score | 57.4 (11.99) | 7.5 (8.7) | 2.2E-04 |
| Current CAPS - intrusion score | 9.8 (1.64) | 0.25 (0.5) | 1.1E-05 |
| Current CAPS - avoidance score | 4.4 (1.52) | 0 (0) | 7.2E-04 |
| Current CAPS - cognition & mood score | 15.4 (4.72) | 2.75 (3.77) | 3.4E-03 |
| Current CAPS - hyperarousal score | 13.2 (2.17) | 2 (3.37) | 5.0E-04 |
| Current CAPS - total severity score | 42.8 (4.91) | 5.0 (7.5) | 7.2E-06 |
| Number of deployments | 3.25 (0.96) | 1.75 (0.5) | 3.2E-02 |
| Shapiro-Wilk test was used to assess normality of continuous variables and either a linear regression or Mann-Whitney U test were implemented accordingly. Discrete variables were examined using a Chi-squared statistic. Abbreviations: BDI, Becks depression inventory; PCL, patient check list, MDD, major depressive disorder; LT, lifetime. |  |  |  |

To confirm the developmental specificity of these data, we performed a large comparative and integrative analysis that included 16 existing transcriptome datasets over 2716 independent samples covering a range of hiPSC-derived neurons and developmentally distinct postmortem brain tissue (see *“Materials and Methods”*). A high degree of transcriptomic convergence was observed between our hiPSC-derived neurons with fetal brain tissue and hiPSC-derived neurons generated from previous reports, confirming their early developmental gene expression profiles (**Figure S4**). Following, the expression variation within our hiPSC-derived neurons were explored. Collectively, known clinical and technical factors explained ~71% of transcriptome variation, with differences between *NGN2*-neurons and mixed FB neurons having the largest genome-wide effect that explained a median ~62% of the observed variation, followed by differences in donor as a repeated measure (~3.2%) (**Figure 1A**). As expected, PCA accurately distinguished mixed FB neurons from *NGN2*-neurons along the first principal component, explaining 78.5% of the variance (**Figure 1B**), with *SOX2* and *TUBB3* as the top genes explaining differences by neuronal cell types (**Figure 1C**). Notably, within each cell type, donor as a repeated measure explained ~28% of transcriptome-wide variance and these genes were strongly enriched for brain eQTLs (**Figure S5**), indicating that the observed inter-individual expression variation reflects genetic regulation of expression. Expression variation due to differing HCort concentrations had a detectable effect on a smaller number of genes.

**Figure 1.**
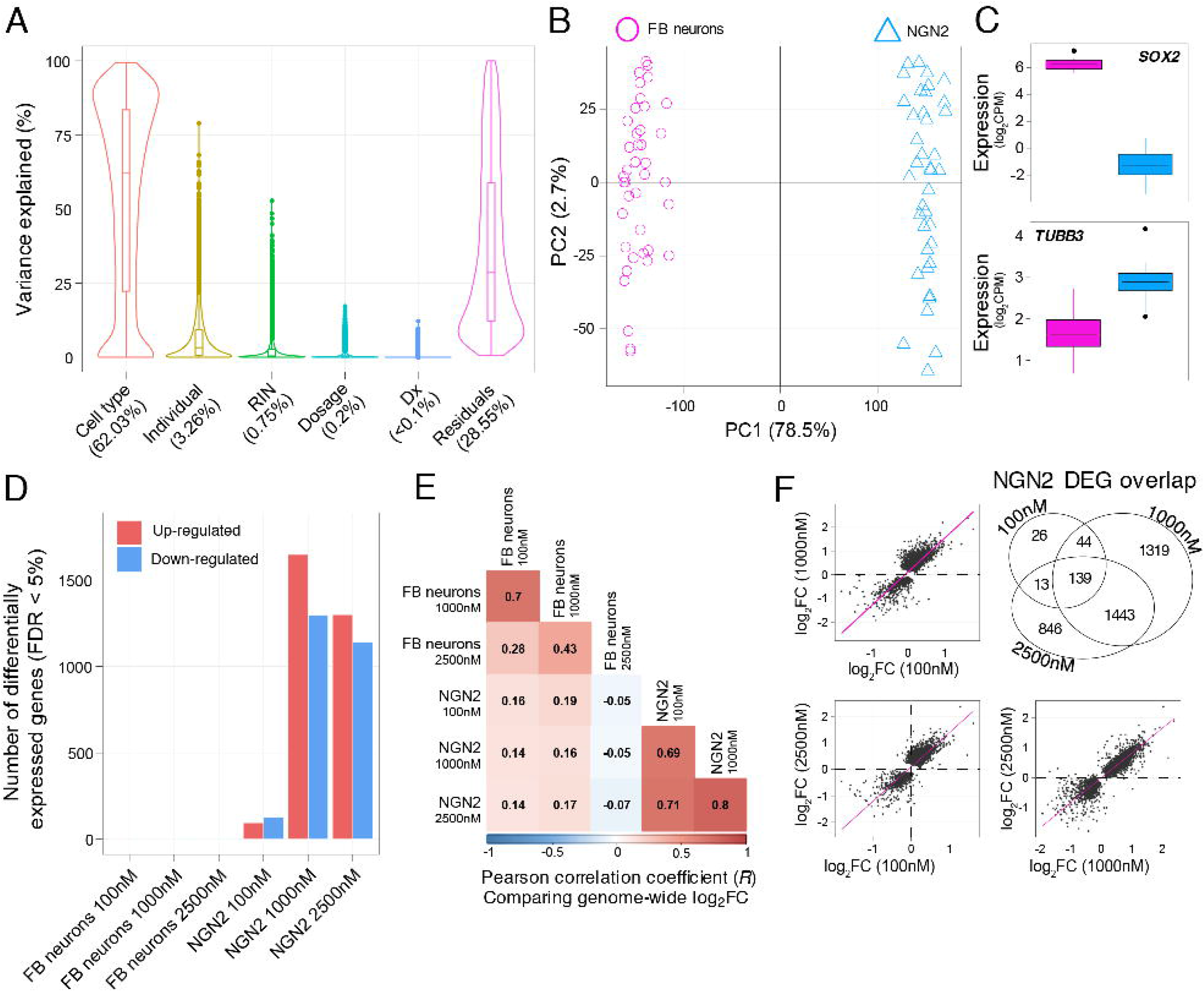
Gene expression changes to HCort in hiPSC-derived neurons. (**A**) To test influence of various factors on gene expression profiles, for each gene, the percentage of gene expression variation attributable to each biological and technical factor was computed. Collectively, these variables explained ~72% of transcriptome variation, with differences between *NGN2*-and mixed forebrain neurons having the largest genome-wide effect that explained a median 62% of the observed variation, followed by differences in donor as a repeated measure (median 3.6%). (**B**) Principal component analysis clearly separates *NGN2*-neurons and mixed forebrain (FB) neurons based on global gene expression. (**C**) An example of two neural genes that show strong cell-type preference. (**D**) Differential gene expression changes in response to increasing concentrations of HCort shows robust changes in *NGN2*-neurons but no changes in mixed FB neurons. (**E**) A comparative analysis of transcriptome-wide log fold-changes in response to different concentrations of HCort across mixed FB and *NGN2*-neurons shows distinct responses across the two cell types. (**F**) A comparative analysis of transcriptome-wide log_2_ fold-changes in response to different concentrations of HCort in *NGN2*-neurons shows similar responses across – indicating a conserved response across all donors to HCort in *NGN2*-neurons.

Dose-dependent transcriptional responses were studied following each concentration of HCort relative to baseline. To focus on the dose-dependent transcriptional responses to HCort and to identify reliable markers of glucocorticoid activation independent of PTSD status, we adjusted for the possible influence of donor as a repeated measure, diagnosis (*e.g*., PTSD+ and PTSD−) and RNA integrity numbers (RIN). Relative to vehicle (0nM HCort), a total of 221, 2,945, and 2,441 genes were significantly differentially expressed (*q*-value< 0.05) in *NGN2*-neurons following incubation with 100 nM, 1000 nM, and 2500 nM of HCort, respectively (**Figure 1D, Supplemental Table 1**). Notably, no single gene was significantly differentially expressed in response to HCort in mixed FB neurons, consistent with a previous report of minimal glucocorticoid response in mixed FB neurons^13^. Transcriptome-wide concordance was examined using HCort-associated log2 fold-changes and divergent patterns of differential responses were observed between mixed FB and *NGN2*-neurons (**Figure 1E**). However, HCort responses within each cell type were highly concordant and largely preserved across increasing concentrations of HCort (**Figure 1E-F**), more specifically in *NGN2*-neurons (**Figure S6**).

To better understand the functional aspects of the transcriptional changes in *NGN2*-neurons, unsupervised weighted gene correlation analysis was applied. Twenty-one co-expression modules (M) were identified (**Figure S7**), and seven were dynamically regulated in response to increasing concentrations of HCort, constituting ~80% of the measured *NGN2*-transcriptome (*n^genes^*=13,571) (**Figure 2, Supplemental Table 2**). For all genes within each module, we compared the absolute effect sizes induced by HCort in *NGN2*-neurons relative to mixed FB neurons (**Figure 2B**); again, HCort treatment elicited significantly enhanced transcriptional response in *NGN2*-neurons relative to mixed FB neurons, which displayed an overall blunted response. Module eigengene (ME) values for M1-4 were all downregulated and enriched for biological processes related to protein translation (M1, *p*=0.002), nucleoside triphosphate processes (M2, *p*=0.003), oxidative phosphorylation (M3, *p*=0.007) and cell cycle (M4, *p*=0.003), respectively. Collectively, these functions have been previously reported to be downregulated with low doses of glucocorticoids across various experimental contexts. Remaining ME values for modules M5-7 all significantly increased with HCort treatment. M5 was significantly up-regulated (*p*=2.1×10^-4^) and strongly enriched for processes related to neurogenesis and cytoskeleton organization, chromatin organization genes, glutamatergic synapses. Likewise, M6 was significantly up-regulated (*p*=3.4×10^-4^) and implicated in trans-synaptic signaling and synapse organization, and ion channel binding. Finally, M7 was up-regulated (*p*=0.009) and enriched for processes related to neurotrophin signaling, MAPKinase signaling, endocrine and inflammatory signaling. Notably, M7 harbored well-known reliable markers of glucocorticoid activation and was significantly enriched for glucocorticoid receptor regulatory genes and harbored a significant fraction of genes with well validated glucocorticoid binding sites in neurons (*p* = 0.003), which are known to have significant glucocorticoid-inducible gene regulatory activity (**Figure S8**).

**Figure 2.**
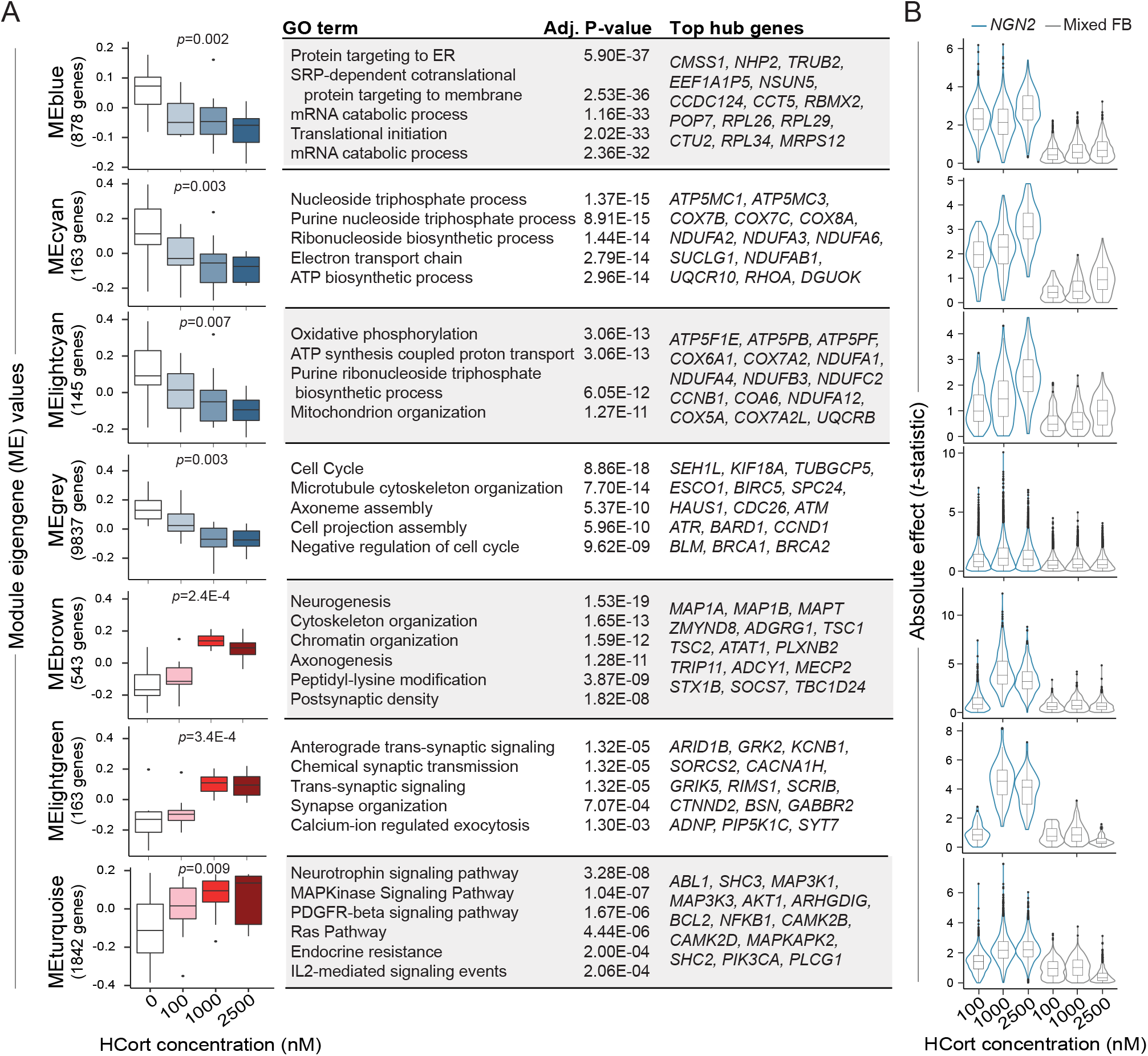
HCort stimulated co-expression modules in *NGN2*-neurons. (**A**) Weighted gene coexpression network analysis (WGCNA) has identify seven groups of co-regulated gene modules. Analysis of variance (ANOVA) was used to assess changes in module eigengene (ME) values with increasing concentration of HCort (*p*-values are labeled above each boxplot). Each module was subjected to gene ontology enrichment analysis and the top most significant enrichment terms and their associated Benjamini-Hochberg adjusted *P*-values are displayed. Further, we also display some of the top hub genes (kME>0.6) within each module for quick interpretation of GR-stimulated gene coexpression modules and candidate individual genes. (**B**) Absolute effect size (using t-statistics from differential gene expression) demonstrate the strength of effect of the various HCort treated conditions in *NGN2*-neurons relative to mixed FB neurons all genes within a module.

Overall, although we observed a weak transcriptional response in mixed FB neurons, HCort treatment of *NGN2*-neurons, independent of diagnosis, resulted in robust down-regulation of protein and nucleic acid metabolism genes, and up-regulation of neurogenesis, chromatin organization genes and synaptic genes.

### PTSD diagnosis-dependent differences in human neurons

Baseline gene expression profiles (vehicle; 0 nM HCort) were compared between PTSD+ and PTSD–groups while adjusting for donor as a repeated measure and the possible influence of RIN (see Materials and methods). No significant differences in gene expression were observed (*q*-value < 0.05) and a distribution of PTSD-related *P*-values, which was no different from the expected uniform distribution, was identified (**Supplemental Table 3**; λ mean = 0.73).

Next, we applied a linear contrast analysis to examine whether genes respond differently to HCort, relative to baseline, in PTSD+ relative to PTSD-combat veterans, here termed differential response genes (DRGs) (**Supplemental Table 4**). We identified 128, 320 and 765 PTSD-specific DRGs in *NGN2*-neurons following 100 nM, 1000 nM and 2500 nM of HCort, respectively, and substantially fewer DRGs in mixed FB neurons (*nominal p*-value < 0.05) (**Figure 3A**). The strongest PTSD-dependent differential response to HCort was observed following 1000 nM of HCort in *NGN2*-neurons (*p*=4.3×10^-32^, linear regression) (**Figure 3B**).

**Figure 3.**
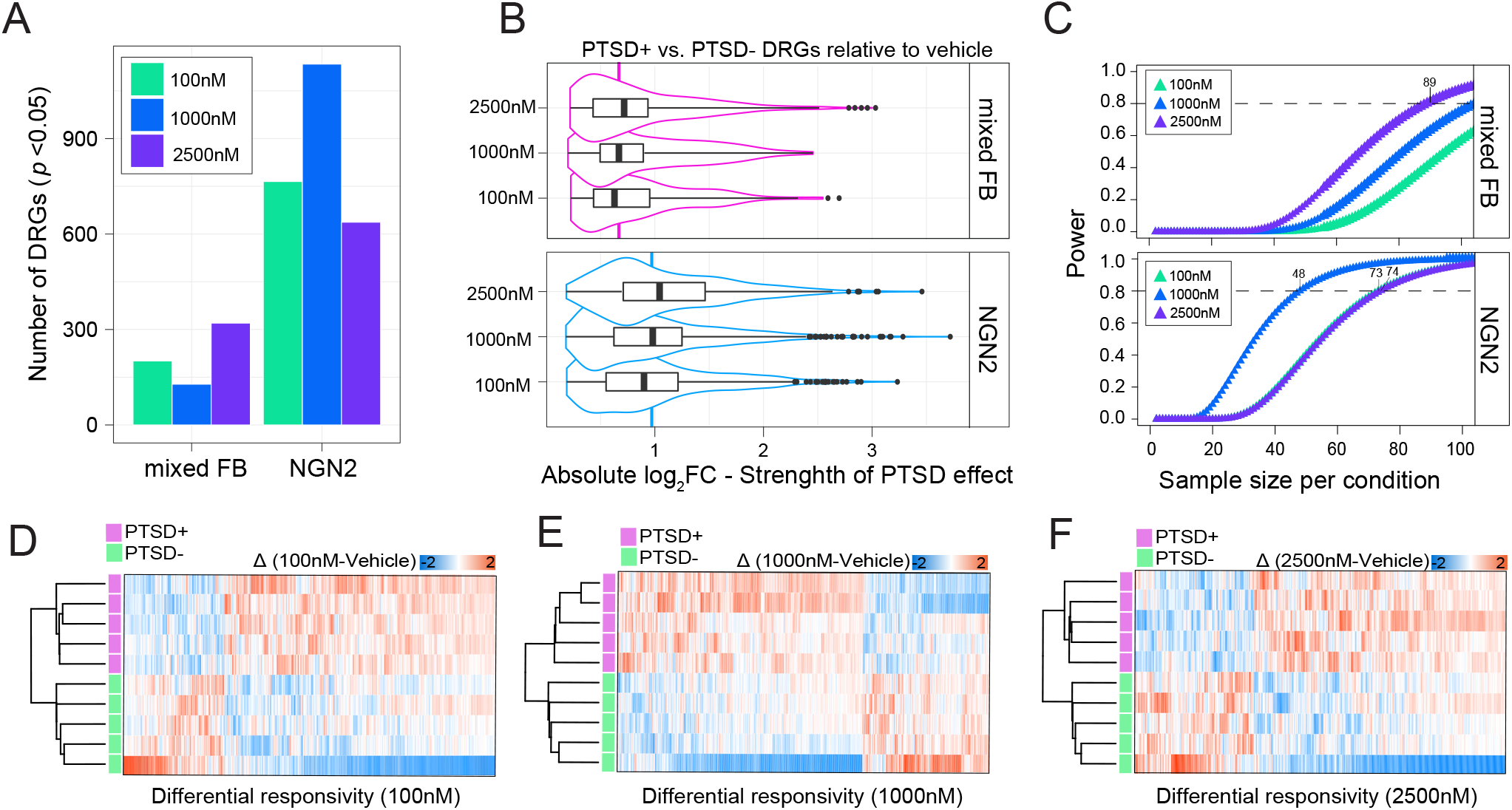
PTSD+ specific responses to HCort in *NGN2*-neurons. (**A**) We conducted an analysis for genes that differ in their response to HCort in PTSD+ donors compared to PTSD-donors, here termed differential response genes (DRGs). Our analysis did not uncover and DRGs that survived multiple test correcting, so we relaxed our P-value significance threshold. (**B**) The absolute log fold-change (logFC) was evaluated for all DRGs passing a nominal p-value significance threshold, p < 0.05. In addition to *NGN2*-neurons showing more DRGs in PTSD+, their strength of effect in PTSD+ was also significantly greater than those identified in mixed forebrain (FB) neurons. (**C**) To inform the design of our future work, we estimated the sample size that is required to identify DRGs specific to PTSD+ that have a power of 0.8 or greater. Our calculation was based on the distribution of nominal *P*-values for each set of DRGs. We set the average probability of detecting an effect to be >0.8 and *α*=0.05. (**D-F**) In the absence of robust PTSD+ effects in neurons, we tested whether the nominally significant *NGN2*-DRGs could correctly classify PTSD+ from PTSD-participants using unsupervised approach.

Given the challenges of low statistical power, substantial intra-donor variation, and the range of complicating factors that can obscure signal, future hiPSC-based studies of PTSD and stress responsivity should be carefully designed to maximize power. To inform the design of future hiPSC-studies of PTSD, we estimated power and sample size for each cell type and HCort dosage using lists of *P*-values derived from DRG analyses. Effect sizes estimated from these data were assumed to be fixed with a nominal error rate *α*=0.05 and several different sample sizes (*n*) were evaluated. In contrast to mixed FB neurons, which required a *n* of 89 (case-control pairs) or greater to reach power of 0.8, the sample size needed for *NGN2*-neurons to reach a power of 0.8 was *n* of 74 for 100 nM, *n* of 48 for 1000 nM and *n* of 74 for 2500 nM (**Figure 3C**). Finally, we explored whether the current set of nominally significant DRGs in *NGN2*-neurons could predict PTSD in this small data set. For each individual, we applied unsupervised classification and observed a clear pattern of HCort dysregulation that correctly classified PTSD+ from PTSD-groups in *NGN2*-neurons (**Figure 3D-F**).

Overall, although we observed robust HCort responses from all ten donors, we resolved relatively weak PTSD-specific HCort responses in *NGN2*-neurons, and recommend that future studies reach sample sizes of at least 50 donor pairs. That being said, our small dataset generated from 5 PTSD cases and 5 controls is sufficient for unsupervised classification of donors by diagnosis based on the HCort response of donor-derived *NGN2*-neurons.

### Replication of PTSD diagnosis-dependent GC-induced changes in PBMCs

Impaired sensitivity of the glucocorticoid receptor to glucocorticoids and increased expression of inflammatory cytokines and innate immune genes are hallmark signatures of PTSD in basal *ex vivo* PBMCs. Our previous work demonstrated the utility of sensitizing PBMCs *in vitro* to the synthetic glucocorticoid dexamethasone (DEX) as a promising method for the exploration of molecular responses to stress hormones in PTSD^9^. Here we sought to confirm these earlier findings by applying identical methods to those previously described to the current cohort of combat-exposed veterans with PTSD (PTSD+; *n*= 5) and without PTSD (PTSD-; *n* = 5) (**Table 1**). In brief, genome-wide RNA-sequencing was generated from cultured PBMCs incubated with three increasing concentrations of DEX (2.5nM, 5nM, 50nM) for all donors in the current study, thereby acting as a truly independent validation set. Following data quality control and pre-processing, we tested the overall response to DEX for each concentration while adjusting for the possible influence of donor as a repeated measure, age, RIN and BMI. Relative to vehicle (0 nM DEX), a total of 0, 1056, and 12,625 genes were significantly differentially expressed (*q*-value < 0.05) following incubation with 2.5 nM, 5 nM, and 50 nM of DEX, respectively (**Figure 4A, Supplemental Table 5**); representing a total of 11,711 unique genes. While the total number of DEX-stimulated genes reported here are lower than those we previously reported, we note that down-sampling our previous study to a comparable sample size (5 PTSD+ vs. 5 PTSD-) captures a similar number of DEX-responsive genes (**Figure 4A**). Moreover, transcriptome-wide concordance of DEX-induced fold changes relative to vehicle between the two independent studies was exceedingly high (average *r*=0.80) (**Figure 4B-C**). Furthermore, we tested whether seven functional gene coexpression modules previously found to be dynamically regulated by DEX were also preserved in their expression profiles in the current dataset using a permutation-based preservation statistic (*Z_summary_*). All DEX-responsive modules displayed strong preservation of their co-expression profiles in the current dataset (**Figure S9**). These findings demonstrate strong conservation of transcriptional changes to DEX in PBMCs that are independent of PTSD and other potentially confounding factors.

**Figure 4.**
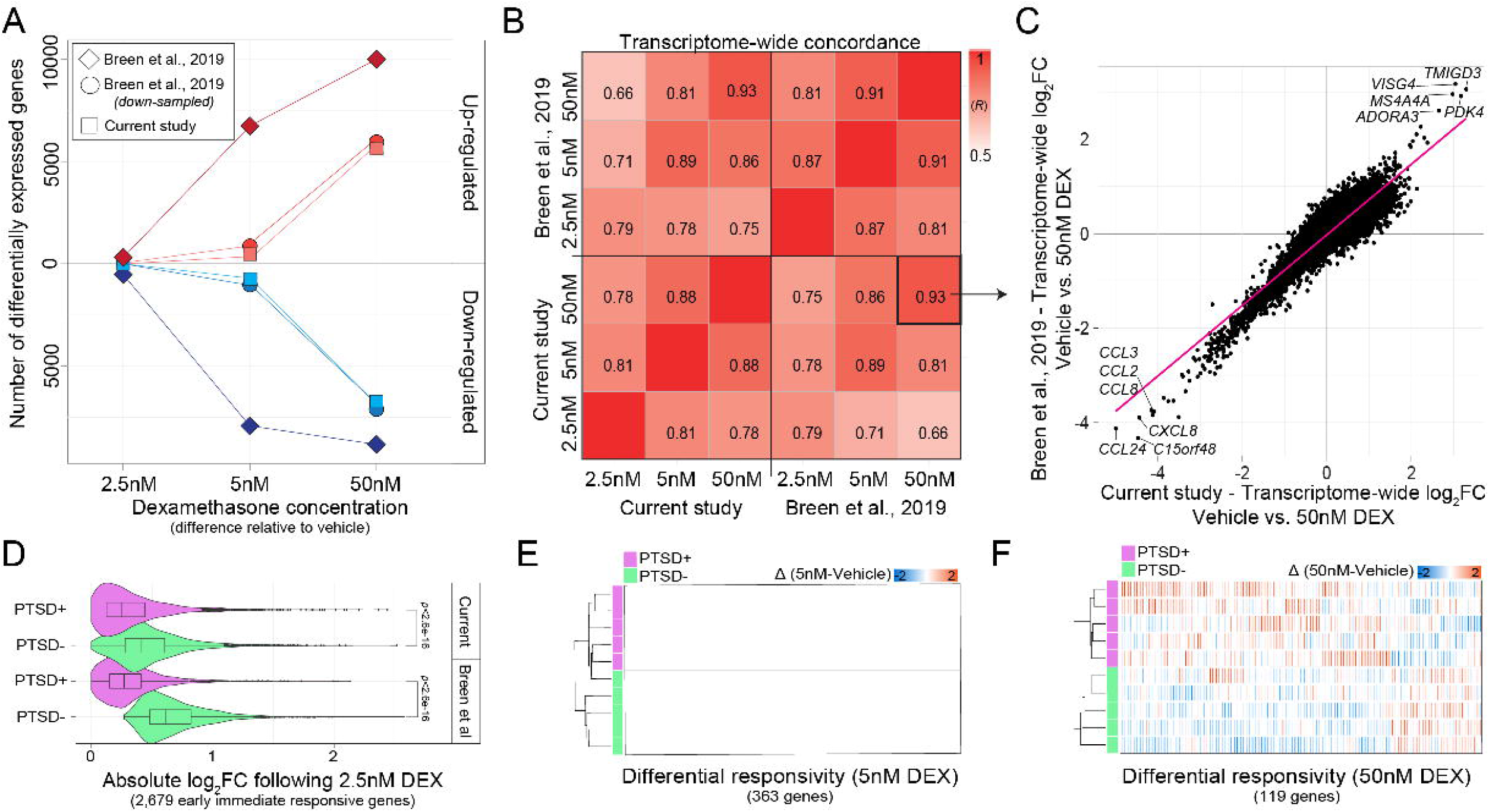
Validation of transcriptional responses to DEX in PBMCs. (**A**) The number of differentially expressed genes observed in the current study (squares) that are up-regulated and down-regulated (y-axis) across three different concentrations of DEX conditioning (x-axis). Note that the current study explored transcriptional responses to DEX in half of the sample size we previously published (Breen *et al*., 2019). Notably, re-analyzed our previous work by down-sampling the total number of individuals to those comparable in the current study and achieved similar numbers of differentially expressed genes. (**B**) Transcriptome-wide log fold-changes in gene expression (differences observed between vehicle and a DEX conditioning) were compared across each DEX concentration and with our previously published work. (**C**) An example of highly conserved transcriptional changes to DEX in the current study (x-axis) and our previous report (y-axis). **(D)** We previously observed that low doses of DEX (2.5 nM) produce robust transcriptional changes in PTSD-participants compared to PTSD+ participants based on 2,679 early immediate responsive genes (Breen *et al*., 2019). Here, we plot the absolute log fold change of these same genes in the current study and reproduce this strong PTSD+ effect. (**E**) We previously observed that mild doses of DEX (5 nM) produce robust transcriptional changes in PTSD+ participants compared to PTSD-participants based on 363 genes (Breen *et al*., 2019). Using these genes, we plot the individual differences (5nM-vehicle) in the current study and reproduce a strong PTSD+ effect. (**F**) We previously observed that mild doses of DEX (5onM) produce robust transcriptional changes in PTSD+ participants compared to PTSD-participants based on 119 genes (Breen *et al*., 2019). Using these genes, we plot the individual differences (50 nM-vehicle) in the current study and reproduce a strong PTSD+ effect.

Our previous work in PBMCs revealed two PTSD diagnosis-dependent effects: i) a robust transcriptional response across 2,679 early immediate responsive genes in PTSD-, but absent in PTSD+ following 2.5 nM DEX; and ii) a differential transcriptional response that was unique to PTSD+ following 5 nM and 50 nM DEX. We queried these early immediate responsive genes following 2.5 nM DEX and confirmed a strong preferential response in PTSD- and an overall blunted response in PTSD+ (Mann-Whitney U test, *p*<2.6e-16) (**Figure 4D**). We also queried whether these genes might also represent expression quantitative trait loci (eQTLs) in the blood and we observed a significant enrichment of eQTLs among these early immediate responsive genes more than expected by random chance (*p*=5.0×10^-9^). We also applied unsupervised classification to test whether genes previously identified to differ in their response to DEX between PTSD+ and PTSD-could accurately predict PTSD+ status in the current independent sample. We observed an 80% and 100% classification accuracy to properly stratify individual-level differences following 5nM and 50nM DEX, respectively, into distinct PTSD+ and PTSD-groups (**Figure 4E-F**). No eQTL enrichment was observed for these two sets of PTSD-specific responses following 5 nM or 50 nM DEX. Overall, these findings confirm our earlier exploratory report, highlighting the utility of our *in vitro* model to capture differences between individuals in sensitivity to GCs and amplify PTSD gene expression effect sizes into a range that may facilitate the development of actionable biomarkers.

Overall, glucocorticoid treatment of PBMCs donor-matched to our hiPSC cohort reproduce glucocorticoid treatment responses observed in our previous PBMC analysis: a robust transcriptional response in PTSD-but not PTSD+ PBMCs to lose dose glucocorticoids (2.5 nM DEX) and a PTSD-specific PBMC response at high dose glucocorticoids (5 nM and 50 nM DEX).

### Comparative analysis of glucocorticoid transcriptional changes in PBMCs and hiPSC neurons

The degree to which stress responses in the blood reflect stress responses in the brain remains poorly resolved. Towards exploring this, we examined the degree of genome-wide concordance of glucocorticoid-induced transcriptional responses between PBMCs and *NGN2*-neurons, observing an overall weak level correlation (*R^2^*=0.02), with many genes and pathways displaying opposite directional effects (**Figure S10**). Next, we studied the degree to which glucocorticoid-responsive gene modules were shared and/or divergent between PBMCs and *NGN2*-derived neurons. This approach applied *Z_summary_* permutation-based preservation statistic (as above), asking whether the underlying gene co-regulatory patterns following glucocorticoid-treatment are preserved across these cellular populations. We identified three glucocorticoid-responsive modules in *NGN2*-neurons that were not preserved in PBMCs, including an up-regulated module implicated in trans-synaptic signaling (*Z_summary_*=1.14) and two down-regulated modules implicated in oxidative phosphorylation (*Z_summary_*=1.68) and nucleoside triphosphate metabolism (*Z_summary_*=0.62) (**Figure 5A-B**). Reciprocally, we identified two (of seven) glucocorticoid-responsive gene co-expression modules identified in PBMCs (described previously) that were not preserved in *NGN2*-neurons, including one up-regulated module implicated in apoptosis (*Z_summary_*=1.31) and one down-regulated module implicated in the inflammatory response and cytokine signaling (*Z_summary_*=1.58) (**Figure 5C**). Both of these modules represent well-known reliable markers of GC activation in PBMCs (*e.g. FKBP5*), harbor a significant fraction of genes with well validated glucocorticoid binding sites (*p* = 0.009, *p* = 0.001, respectively), and display clear dose-response effects that were absent in *NGN2*-neurons (**Figure 5D**). Notably these dose response genes do not show a strong enrichment of cis-eQTLs, and thus are not thought to be genetically regulated and vary between individuals (**Figure S5**). Taken together, these results suggest that glucocorticoid-responsive pathways and processes are largely distinct between *NGN2*-neurons and PBMCs, and highlight several fundamental glucocorticoid responses that are unique to either tissue/cell type.

**Figure 5.**
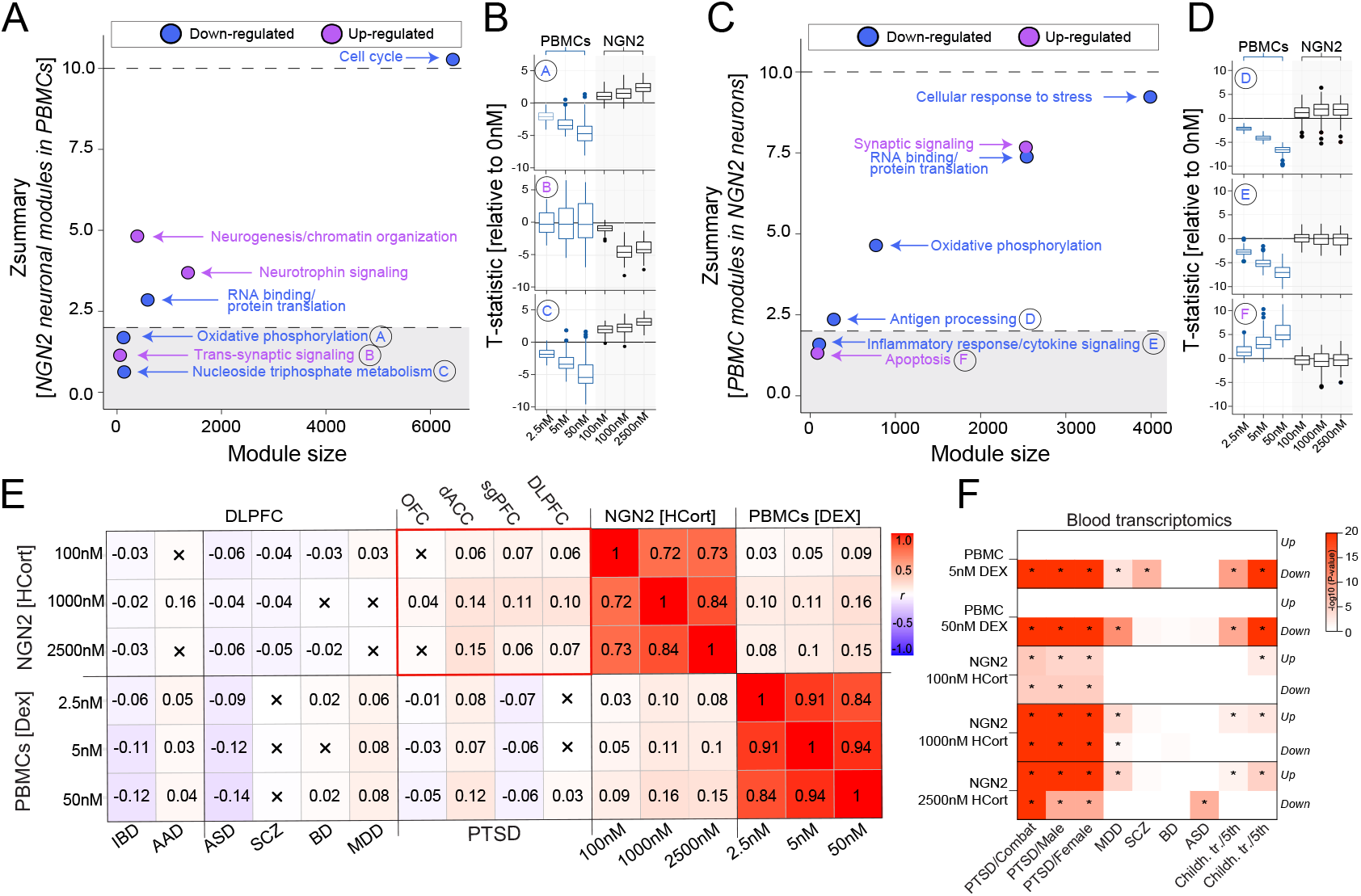
Concordance of glucocorticoid-stimulated gene expression between neurons and PBMCs. Gene set preservation analysis identifies GC-responsive modules that are shared and unique across PBMCs and *NGN2*-neurons. (**A**) Two GC-responsive modules in PBMCs displayed no preservation (Z_summary_ < 2) in *NGN2*-neurons and (**B**) the global effect size, relative to an untreated condition, was significantly greater in PBMCs relative to *NGN2*-neurons. In the reverse approach, (**C**) three GC-responsive modules in *NGN2*-neurons displayed no preservation (Z_summary_ < 2) in PBMCs and (**D**) the global effect size, relative to an untreated condition, was significantly greater in *NGN2*-neurons relative to PBMCs. **(E)** We compared transcriptome-wide changes in gene expression following HCort exposure in *NGN2-and* mixed FB neurons with expression changes following DEX exposure in PBMCs. We observed significant positive correlations in expression changes between *NGN2*-neurons and PBMCs and significant negative correlations in expression between mixed FB neurons and PBMCs. (**F**) Convergence of GC-induced transcriptional responses with findings from large-scale blood transcriptomic studies using a Fisher’s exact test.

### Concordance between glucocorticoid transcriptional responses in vitro and transcriptional dysregulation in patient postmortem brain tissues and ex vivo PBMCs

Insights into how stress responses in blood and brain cells echo transcriptional changes observed in the brain of individuals with PTSD and related neuropsychiatric disorders remain limited. Comparison of differential gene expression log_2_ fold change signatures revealed a positive, significant overlap between transcriptional responses to HCort in *NGN2*-neurons and transcriptional dysregulation observed across three postmortem brain regions from individuals with PTSD (**Figure 5E**). This level of genome-wide concordance was substantially lower when contrasting HCort responses in *NGN2*-neurons with transcriptional dysregulation in major depressive disorder (MDD), bipolar disorder (BD), schizophrenia (SCZ) and autism spectrum disorder (ASD). This level of transcriptional convergence was not observed when comparing DEX-induced transcriptional responses in PBMCs with these postmortem brain tissue summary statistics. Notably, we also observed that a significant fraction of genes that respond to HCort in *NGN2*-neurons implicate genetic risk loci for neurodevelopmental disorders (**Figure S11**), consistent with recent reports (ref).

We reciprocally tested whether glucocorticoid-responsive genes in either PBMCs or *NGN2*-neurons were enriched for transcriptional dysregulation observed in PBMCs from individuals with PTSD and other neuropsychiatric disorders. A significant overlap was observed between PTSD-related transcriptomic dysregulation in the blood as a consequence of combat trauma and interpersonal traumas with DEX-induced signatures in PBMCs and HCort-induced signatures in *NGN2*-neurons (**Figure 5F**). A significant convergence of gene dysregulation was also observed between glucocorticoid-induced transcriptional responses in both PBMCs and *NGN2*-neurons with childhood trauma-related transcriptomic dysregulation in the blood. Notably, we did not observe substantial convergence with schizophrenia or BD-related transcriptomic changes in the blood. Collectively, these results suggest that GC-induced transcriptional responses reflect both PTSD-related transcriptional responses across both blood and brain cells.

Overall, we observe largely distinct glucocorticoid responses between *NGN2*-neurons and PBMCs, with neuron-specific responses enriched for synaptic genes, and PBMC-specific responses for inflammatory response genes. Neuronal, but not PBMC, HCort responses showed a positive and significant overlap with three postmortem brain tissue samples from individuals with PTSD.

## DISCUSSION

Here we applied an hiPSC-based model to explore the cell-type-specific and diagnosis-dependent responses of blood and brain cells derived from PTSD cases and combat-exposed controls to glucocorticoid treatment. First, we observe largely distinct glucocorticoid responses between *NGN2*-neurons and PBMCs, with neuron-specific responses enriched for synaptic genes, and PBMC-specific responses for inflammatory response genes. Only neuronal HCort responses showed a positive and significant overlap with postmortem brain tissue from patients with PTSD. Second, our results highlight the need for larger and better-powered hiPSC-based studies of complex genetic psychiatric disorders, consistent with other reports^28^. Although the small PTSD-specific HCort responses observed here were sufficient for unsupervised classification of donors by diagnosis based on the HCort response of donor-derived *NGN2*-neurons, we recommend that future studies reach sample sizes of at least 50 donor pairs. The work provides a proof-of-principle demonstration that disease-relevant processes not detectable at baseline can emerge in more physiologically relevant contexts (*e.g*. glucocorticoid response).

Much work still remains in order to resolve the cell type(s) of origin in PTSD. Are blood-specific glucocorticoid changes a critical part of disease risk, resulting in the release of additional cytokines and inflammatory molecules that impact brain function? Or are they simply an incomplete biomarker of events occurring in the brain? Within the brain, are neuron-specific responses adequate to interpret G×E interactions? Certainly future experiments should investigate a larger variety of neuronal and glial cell fates. While it is true that we observed an unexpectedly weak transcriptional response in mixed FB neuronal populations that include both neurons and astrocytes, this does not rule out a critical role for glial cells in PTSD etiology, as the differences observed here could instead reflect compensatory effects mediated by astrocytes that weakened the observed neuronal HCort-response, and/or a reduction in statistical power owing to cellular heterogeneity and variation in cell type composition between donors and experiments in mixed FB neurons. Because robust and distinct cell-type-specific glucocorticoid responses were observed in radial glia, neural progenitors, and neurons in the context of 3D brain organoids by single cell RNAseq^15^, we suggest our findings likely reflect the latter.

It is important to consider that stress impacts risk to psychiatric disorders throughout the lifespan, across prenatal development, childhood, adolescence, as well as in adulthood. As described above, hiPSC-derived neurons most closely resemble fetal brain cells^33–36^. Consequently, our hiPSC-based model may in fact be capturing the G×E impact of fetal stress exposure on PTSD risk. Even here, the biological relevance is high, as increased glucocorticoid signaling during fetal development has been linked to psychiatric disorders^37,38^. Towards this, glucocorticoid treatment increases the proliferation of hiPSC-derived neural progenitor cells^39^, and impairs neuronal differentiation and maturation in hiPSC-derived neurons^15,40^ and primary mouse neurons^41^. It follows from these observations that if hiPSC-derived neurons represent a fetal-like pre-trauma state, then patient-derived PBMCs represent an adult state, post-trauma and disease onset. Thus, we cannot rule-out that the cell-type-specific effects described here are in fact maturity-dependent, trauma-dependent, and/or PTSD illness etiology-dependent. Nonetheless, hiPSC-derived neurons represent a novel and exciting platform with which to screen for genetic, epigenetic, and/or pharmacological interventions that can prevent, ameliorate, or reverse acute or chronic neuronal responses to stress. We acknowledge that hiPSC modelling of epigenetic factors is challenging due to the resting of the epigenomic landscape during hiPSC reprogramming.

The mechanisms by which the growing list of genetic loci associated with PTSD impact disease pathophysiology are unknown. In particular, interactions between these myriad variants and the environment are undetermined, limiting the translation of complex genetic insights into medically actionable information. Despite substantial advances in explaining brain disease risk through annotation of GWAS signals with functional genomics, these approaches have focused exclusively on genetically regulated gene expression and genetic regulation of higher order biology, without accounting for the impact of environment or stressors. This failure to incorporate G×E interactions is important across studies of neuropsychiatric disorders, many of which require or include specific stressors (e.g. PTSD) or exposures (e.g. substance use disorders, anorexia nervosa) among diagnostic criteria, or count exposure to illicit substances or extreme stress as among the most critical risk factors for disease onset and development (e.g. schizophrenia, anorexia nervosa, psychotic disorders more broadly). Thus, given the critical importance of gene x stress (and G×E more broadly), we believe that moving forward, functional genomics approaches must integrate stem cell models and genome engineering, in order to resolve the impact of patient-specific variants across cell types and environmental stressors. Our hope is that dissecting how disease risk variants interact with the environment across the diverse cell types populating the human brain, will improve diagnostics, predict clinical trajectories, and identify pathways that could serve as pre-symptomatic points of therapeutic intervention.

## Acknowledgements

This work was supported by the Office of the Assistant Secretary of Defense for Health Affairs through the USAMRMC BAA for Extramural Medical Research under Award No. W81XWH-15-1-0706. Opinions, interpretations, conclusions, and recommendations are those of the author and are not necessarily endorsed by the Department of Defense. This material is the result of work supported with resources and the use of facilities at the James J. Peters VAMC in Bronx, NY. The contents of this manuscript do not represent the views of the U.S. Department of Veterans Affairs or the United States Government. Further, this work was supported by the New York Stem Cell Foundation Research Institute (NYSCF).

## Author contributions

Study conception was provided by R.Y. and K.B. Neuronal generation and modeling was provided M.C. and T.R. Data analysis was provided by M.S.B., C.S. and C.X. All authors interrelated data and critically revised the manuscript for important intellectual content. Data and specimen collections were carried out by H.B., J.D.F., L.M.B. and R.Y. All authors wrote and approved the final manuscript.

## Data availability

All raw RNA-seq FASTQ files have been uploaded to the gene expression omnibus under accession number XXXXXXX (will release following manuscript publication).

## Code availability

All computational code is available at GitHub: https://github.com/BreenMS

## Supplemental Author List for NYSCF Global Array Team

Lauren Bauer, Katie Brenner, Geoff Buckley-Herd, Sean DesMarteau, Patrick Fenton, Peter Ferrarotto, Jordan Goldberg, Jenna Hall, Selwyn Jacob, Travis Kroeker, Greg Lallos, Hector Martinez, Paul McCoy, Rick Monsma, Dorota Moroziewicz, Reid Otto, Katie Reggio, Bruce Sun, Rebecca Tibbets, Dong Woo Shin, Monica Zhou, Matthew Zimmer

## REFERENCES

1. Kessler, R. C. et al. Prevalence and treatment of mental disorders, 1990 to 2003. N Engl J Med 352, 2515–2523, doi:10.1056/NEJMsa043266 (2005).

2. Kremen, W. S., Koenen, K. C., Afari, N. & Lyons, M. J. Twin studies of posttraumatic stress disorder: differentiating vulnerability factors from sequelae. Neuropharmacology 62, 647–653, doi:10.1016/j.neuropharm.2011.03.012 (2012).

3. Wolf, E. J. et al. A classical twin study of PTSD symptoms and resilience: Evidence for a single spectrum of vulnerability to traumatic stress. Depress Anxiety 35, 132–139, doi:10.1002/da.22712 (2018).

4. Nievergelt, C. M. et al. International meta-analysis of PTSD genome-wide association studies identifies sex-and ancestry-specific genetic risk loci. Nat Commun 10, 4558, doi:10.1038/s41467-019-12576-w (2019).

5. Duncan, L. E. et al. Largest GWAS of PTSD (N=20 070) yields genetic overlap with schizophrenia and sex differences in heritability. Mol Psychiatry 23, 666–673, doi:10.1038/mp.2017.77 (2018).

6. Gelernter, J. et al. Genome-wide association study of post-traumatic stress disorder reexperiencing symptoms in >165,000 US veterans. Nat Neurosci 22, 1394–1401, doi:10.1038/s41593-019-0447-7 (2019).

7. Girgenti, Matthew J., et al. “Transcriptomic organization of the human brain in post-traumatic stress disorder.”Nature Neuroscience 24.1 (2021): 24–33.

8. Yehuda, R., Golier, J. A., Yang, R. K. & Tischler, L. Enhanced sensitivity to glucocorticoids in peripheral mononuclear leukocytes in posttraumatic stress disorder. Biol Psychiatry 55, 1110–1116, doi:10.1016/j.biopsych.2004.02.010 (2004).

9. Breen, M. S. et al. Differential transcriptional response following glucocorticoid activation in cultured blood immune cells: a novel approach to PTSD biomarker development. Translational psychiatry 9, 201, doi:10.1038/s41398-019-0539-x (2019).

10. Daskalakis, N. P., Cohen, H., Cai, G., Buxbaum, J. D. & Yehuda, R. Expression profiling associates blood and brain glucocorticoid receptor signaling with trauma-related individual differences in both sexes. Proc Natl Acad Sci U S A 111, 13529–13534, doi:10.1073/pnas.1401660111 (2014).

11. Yehuda, R. et al. Lower methylation of glucocorticoid receptor gene promoter 1F in peripheral blood of veterans with posttraumatic stress disorder. Biol Psychiatry 77, 356–364, doi:10.1016/j.biopsych.2014.02.006 (2015).

12. Cathomas, F., Murrough, J. W., Nestler, E. J., Han, M. H. & Russo, S. J. Neurobiology of Resilience: Interface Between Mind and Body. Biol Psychiatry 86, 410–420, doi:10.1016/j.biopsych.2019.04.011 (2019).

13. Lorsch, Z. S. et al. Stress resilience is promoted by a Zfp189-driven transcriptional network in prefrontal cortex. Nat Neurosci 22, 1413–1423, doi:10.1038/s41593-019-0462-8 (2019).

14. Lieberman, R., Kranzler, H. R., Levine, E. S. & Covault, J. Examining FKBP5 mRNA expression in human iPSC-derived neural cells. Psychiatry Res 247, 172–181, doi:10.1016/j.psychres.2016.11.027 (2017).

15. Cruceanu, C. et al. Cell-type specific impact of glucocorticoid receptor activation on the developing brain. bioRxiv, 2020.2001.2009.897868, doi:10.1101/2020.01.09.897868 (2020).

16. Brennand, K. et al. Phenotypic differences in hiPSC NPCs derived from patients with schizophrenia. Mol Psychiatry 20, 361–368, doi:10.1038/mp.2014.22 (2015).

17. Paull, D., Sevilla, A., Zhou, H. et al. Automated, high-throughput derivation, characterization and differentiation of induced pluripotent stem cells. Nat Methods 12, 885–892, doi.org/10.1038/nmeth.3507 (2015).

18. Kahler D.J. et al. Improved methods for reprogramming human dermal fibroblasts using fluorescence activated cell sorting. PLoS One 8, e59867. doi: 10.1371/journal.pone.0059867 (2013)

19. Topol, A., Tran, N. N. & Brennand, K. J. A guide to generating and using hiPSC derived NPCs for the study of neurological diseases. Journal of visualized experiments: JoVE, e52495, doi:10.3791/52495 (2015).

20. Bowles, K. R., T, C. W. J., Qian, L., Jadow, B. M. & Goate, A. M. Reduced variability of neural progenitor cells and improved purity of neuronal cultures using magnetic activated cell sorting. PLoS One 14, e0213374, doi:10.1371/journal.pone.0213374 (2019).

21. Zhang, Y. et al. Rapid single-step induction of functional neurons from human pluripotent stem cells. Neuron 78, 785–798. doi.org/10.1016/j.neuron.2013.05.029 (2013)

22. Bolger, A. M., Lohse, M. & Usadel, B. Trimmomatic: a flexible trimmer for Illumina sequence data. Bioinformatics30, 2114–2120 (2014).Return

23. Dobin, A. et al. STAR: ultrafast universal RNA-seq aligner. Bioinformatics 29, (15–21 (2013).

24. Liao, Y., Smyth, G. K. & Shi, W. featureCounts: an efficient general purpose program for assigning sequence reads to genomic features. Bioinformatics 30, 923–930 (2013).

25. Breen, Michael S., et al. “Transcriptional signatures of participant-derived neural progenitor cells and neurons implicate altered Wnt signaling in Phelan-McDermid syndrome and autism.”Molecular autism 11.1 (2020): 1–23.

26. Hoffman, G. E. et al. Transcriptional signatures of schizophrenia in hiPSC-derived NPCs and neurons are concordant with post-mortem adult brains. Nat Commun 8, 2225, doi:10.1038/s41467-017-02330-5 (2017).

27. Ritchie, M. E. et al. limma powers differential expression analyses for RNA-sequencing and microarray studies. Nucleic Acids Res.43, e47–e47 (2015).

28. Hoffman, Gabriel E., and Eric E. Schadt. “variancePartition: interpreting drivers of variation in complex gene expression studies.”BMC bioinformatics 17.1 (2016): 1–13.

29. Zhang, B. & Horvath, S. A general framework for weighted gene co-expression network analysis. Stat Appl Genet Mol Biol.4, 17 (2005).Return

30. Chen, J., Bardes, E. E., Aronow, B. J. & Jegga, A. G. ToppGene Suite for gene list enrichment analysis and candidate gene prioritization. Nucleic Acids Res.37(suppl_2), W305–W311 (2009).

31. Langfelder, Peter, et al. “Is my network module preserved and reproducible?” PLoS computational biology 7.1 (2011): e1001057.

32. Megan Orr and Peng Liu, The R Journal (2009) 1:1, pages 47–53.

33. Mariani, J. et al. Modeling human cortical development in vitro using induced pluripotent stem cells. Proc Natl Acad Sci U S A 109, 12770–12775, doi:1202944109 [pii] 10.1073/pnas.1202944109 (2012).

34. Pasca, A. M. et al. Functional cortical neurons and astrocytes from human pluripotent stem cells in 3D culture. Nat Methods 12, 671–678, doi:10.1038/nmeth.3415 (2015).

35. Qian, X. et al. Brain-Region-Specific Organoids Using Mini-bioreactors for Modeling ZIKV Exposure. Cell 165, 1238–1254, doi:10.1016/j.cell.2016.04.032 (2016).

36. Nicholas, C. R. et al. Functional maturation of hPSC-derived forebrain interneurons requires an extended timeline and mimics human neural development. Cell stem cell 12, 573–586, doi:S1934-5909(13)00141-0 [pii] 10.1016/j.stem.2013.04.005 (2013).

37. Carson, R., Monaghan-Nichols, A. P., DeFranco, D. B. & Rudine, A. C. Effects of antenatal glucocorticoids on the developing brain. Steroids 114, 25–32, doi:10.1016/j.steroids.2016.05.012 (2016).

38. Buss, C. et al. Intergenerational Transmission of Maternal Childhood Maltreatment Exposure: Implications for Fetal Brain Development. J Am Acad Child Adolesc Psychiatry 56, 373–382, doi:10.1016/j.jaac.2017.03.001 (2017).

39. Ninomiya, E. et al. Glucocorticoids promote neural progenitor cell proliferation derived from human induced pluripotent stem cells. Springerplus 3, 527, doi:10.1186/2193-1801-3-527 (2014).

40. Raciti, M. et al. Glucocorticoids alter neuronal differentiation of human neuroepithelial-like cells by inducing long-lasting changes in the reactive oxygen species balance. Neuropharmacology 107, 422–431, doi:10.1016/j.neuropharm.2016.03.022 (2016).

41. Provencal, N. et al. Glucocorticoid exposure during hippocampal neurogenesis primes future stress response by inducing changes in DNA methylation. Proc Natl Acad Sci U S A 117, 23280–23285, doi:10.1073/pnas.1820842116 (2020).

